# Effects of anti-fibrotic standard of care drugs on senescent human lung fibroblasts

**DOI:** 10.1101/2022.05.06.490939

**Authors:** Stephanie B. Garcia, Miriam S. Hohmann, Ana Lucia Coelho, Waldiceu A. Verri, Cory M. Hogaboam

## Abstract

**Rationale:** Cellular senescence is crucial in the progression of idiopathic pulmonary fibrosis (IPF), but it is yet unclear whether the standard-of-care (SOC) drugs nintedanib and pirfenidone have senolytic properties.

**Objectives:** We attempted to illuminate the effects of SOC drugs on senescent normal and IPF lung fibroblasts *in vitro*.

**Methods:** Colorimetric/fluorimetric assays, qRT-PCR, and western blotting were used to evaluate the effect of SOC drugs on senescent normal and IPF lung fibroblasts.

**Results:** SOC drugs did not induce apoptosis in the absence of death ligands in either normal or IPF senescent cells. Nintedanib increased caspase-3 activity in the presence of Fas Ligand (FasL) in normal but not in IPF senescent fibroblasts. Conversely, nintedanib enhanced B cell lymphoma (Bcl)-2 expression in senescent IPF lung fibroblasts. Moreover, in senescent IPF cells, pirfenidone alone induced mixed lineage kinase domain-like pseudokinase (MLKL) phosphorylation, provoking necroptosis. However, fragmented gasdermin D, indicating pyroptosis, was not detected under any condition. In addition, SOC drugs increased transcript levels of fibrotic and senescence markers in senescent IPF fibroblasts, whereas D+Q inhibited all these markers. Finally, D+Q enhanced growth differentiation factor 15 (GDF15) transcript and protein levels in both normal and IPF senescent fibroblasts.

**Conclusions:** In the presence and absence of the extrinsic pro-apoptotic ligands, SOC drugs failed to trigger apoptosis in senescent fibroblasts, possibly due to enhanced Bcl-2 levels and the activation of the necroptosis pathway. SOC drugs elevated fibrotic and senescence markers in IPF lung fibroblasts. Together, these data demonstrated the inefficacy of SOC in targeting senescent cells. Further investigation is required to fully elucidate the therapeutic implications of SOC drugs on other senescent cell types in IPF.

## Introduction

Idiopathic pulmonary fibrosis (IPF) is a chronic, progressive, and the most common type of idiopathic interstitial lung disease, characterized by histopathologic and/or radiologic findings of usual interstitial pneumonia (MARTINEZ; COLLARD; PARDO; RAGHU *et al*., 2017; RAGHU; CHEN; HOU; YEH *et al*., 2016). IPF is an age-related disease that occurs in individuals older than 50 years, and its median survival is approximately 3 to 5 years after diagnosis (RAGHU; CHEN; HOU; YEH *et al*., 2016). The disease course is heterogeneous and includes dyspnea, worsening lung function, and impaired quality of life (KING; PARDO; SELMAN, 2011; MAHER; KREUTER; LEDERER; BROWN *et al*., 2019).

Several risk factors have been described to enhance the risk of IPF, including genetics, gender, aging, comorbidities, smoking, environmental and occupation exposure. Nonetheless, aging is the most prominent factor, and recent studies have highlighted the contribution of senescence cells in IPF (American Thoracic Society. Idiopathic pulmonary fibrosis: diagnosis and treatment. International consensus statement. American Thoracic Society (ATS), and the European Respiratory Society (ERS), 2000). Senescent cells secrete an array of chemokines, pro-inflammatory cytokines, growth factors, and extracellular matrix proteases, thus comprising the senescence-associated secretory phenotype (SASP) (SALAMA; SADAIE; HOARE; NARITA, 2014).

Lung fibroblasts are essential in wound healing in response to lung injury (KENDALL; FEGHALI-BOSTWICK, 2014). When the lung epithelium is damaged, activated fibroblasts differentiate in myofibroblasts and migrate to the injury site, producing several extracellular matrix (ECM) components to promote tissue repair (PARIMON; HOHMANN; YAO, 2021). On the one hand, accumulated fibroblasts become senescent and reduce their ECM deposition, limiting the fibrosis process. On the other hand, growing evidence has shown that targeting senescent fibroblasts reduces pulmonary fibrosis in mice models. In this sense, senescent fibroblasts have been associated with the pathogenesis of IPF.

Senescence cells play a deleterious role in IPF, and it has been shown that the removal of senescence cells increases the life span of animal models (BAKER; CHILDS; DURIK; WIJERS *et al*., 2016). However, the mechanism how senescence cells exacerbate the disease remains unclear. Although there are many cellular senescence hallmarks described, the senescence phenotype is diverse, and its mechanisms are not conserved, which makes it a challenge in terms to finding a specific marker (SOTO-GAMEZ; QUAX; DEMARIA, 2019).

The senescence-associated β-galactosidase (SA-βgal) was the first marker described as a potent identification of senescent cells. However, proliferating cells also express β-galactosidase, which makes it a non-reliable marker. Researchers rely on checking the presence of two to four markers to confirm the presence of SC, such as the expression of senescence-associated β-galactosidase (CAMPISI; D’ADDA DI FAGAGNA, 2007), Cyclin-Dependent Kinase Inhibitor (CDKN)2A (p16), CDKN1A (p21), besides the expression of key SASP factors like Interleukin (IL)-6 or IL-1α (FAGET; REN; STEWART, 2019), Wnt16 (BINET; YTHIER; ROBLES; COLLADO *et al*., 2009) and more recently GDF15 (BASISTY; KALE; JEON; KUEHNEMANN *et al*., 2020). That stated, the identification of mechanisms to remove senescent cells would have a remarkable impact on the quality of life and burden of IPF.

For years, the only treatment available for IPF was transplantation. However, in 2014, The United States Food and Drug Administration (FDA) approved nintedanib (Ofev, Boehringer Ingelheim) and pirfenidone (Esbriet, InterMune) for the treatment of this disease (KARIMI-SHAH; CHOWDHURY, 2015). Nintedanib, a tyrosine kinase inhibitor, blocks the effects of platelet-derived growth factor, fibroblast growth factor, and vascular endothelial growth factor receptor, growth factors that play a role in IPF (RAGHU; SELMAN, 2015). Moreover, nintedanib inhibits signaling pathways in the proliferation, migration, and maturation of lung fibroblasts (HOSTETTLER; ZHONG; PAPAKONSTANTINOU; KARAKIULAKIS *et al*., 2014; RAGHU; SELMAN, 2015). Whereas pirfenidone, whose mechanism remains unclear, exerts anti-fibrotic, antioxidant, and anti-inflammatory effects to reduce lung collagen synthesis and deposition in bleomycin animal models. Although the FDA approval of nintedanib and pirfenidone, the standard-of-care (SOC) drugs, expresses a significant advance by offering the first available options for IPF patients, such therapies do not cure IPF or significantly improve the quality of life of IPF patients (KATO; SHIN; PALUMBO; PAPAGEORGIOU *et al*., 2021).

Senolytics are drugs that can selectively induce senescent cell apoptosis (KELLOGG; KELLOGG; MUSI; NAMBIAR, 2021). The cocktail Dasatinib plus Quercetin (D+Q) constitutes the first combination of senolytics described. The combination of a tyrosine kinase inhibitor and a flavonoid with antioxidant properties, respectively, ameliorates bleomycin-induced pulmonary fibrosis and improves pulmonary and physical function and body composition (SCHAFER; WHITE; IIJIMA; HAAK *et al*., 2017). In addition, D+Q is described to provoke apoptosis of senescent cells in human tissue and alleviate several age-related disorders in mice (JUSTICE; NAMBIAR; TCHKONIA; LEBRASSEUR *et al*., 2019).

Bearing in mind that therapeutic approaches that diminish senescent cells tend to attenuate the progression of pulmonary fibrosis, we compared the effects of SOC drugs and D+Q on apoptosis-resistant senescent IPF fibroblasts. In addition, we evaluate the cell death types that could be involved after SOC drugs treatment.

## Materials and Methods

### Study approval

This Institutional Review Board at Cedars-Sinai Medical Center approved all experiments with primary human tissue, and informed consent was obtained before inclusion in the studies described.

### Senescent fibroblast generation

Primary normal lung fibroblasts were derived from nonfibrotic lung samples without signs of disease from lung explants. Senescent fibroblasts were cultured in Dulbecco’s Modified Eagle Medium (DMEM; Lonza, Basel, Switzerland) supplemented with 15% fetal bovine serum (FBS; Atlas Biologicals, Inc, Fort Collins, CO), 1% penicillin/streptomycin (Mediatech, Manassas, VA), 1% glutamine (Mediatech) and 0.1% of primocin (InvivoGen, San Diego, CA) at 37 C, and 10% CO_2_. To obtain senescent fibroblasts, proliferative normal and IPF lung fibroblasts were repeatedly passaged in culture until they reached a senescent morphological phenotype (enlarged, flattened, and irregular shape) and SA-βgal activity (HOHMANN; HABIEL; COELHO; VERRI *et al*., 2019).

### Cell viability

Cell viability was evaluated using AlamarBlue Cell Viability Reagent (ThermoFisher Scientific, Waltham, MA, USA). Senescent lung fibroblasts (3 × 10^4^ cells/well) were treated with nintedanib (Ofev^®^, Boehringer Ingelheim, Germany; 300nM), pirfenidone (Esbriet^®^, Genentech, San Francisco; 2.5 mM) or dasatinib (Tocris, Bristol, UK; 20μM) + quercetin (Sigma-Aldrich, St. Louis, MO; 15μM) for 24 hours (h) followed by 3h of Super FasL (100ng/ml). AlamarBlue Cell Viability Reagent was added to the cells and incubated for 4 hours at 37°C. Fluorescence values were assessed using a fluorescence excitation wavelength of 560 nm and an emission of 590 nm.

### Caspase-3 assay

The effect of SOC drugs on caspase-3 activity was observed by using Caspase-3/CPP32 Fluorometric Assay Kit (BioVision Inc., Milpitas, CA, USA). Senescent fibroblasts were treated with SOC drugs or D+Q for 24h, followed by 3h of Super FasL stimuli. Cells were lysed in 50 μl chilled cell lysis buffer on ice for 10 min before 50 μl of 2X reaction buffer (containing 10 mM Dithiothreitol) was added, followed by 50 μM DEVD-AFC substrate, inc were lysed in Trizol ubated at 37 °C for 2 h. Fluorescence was measured at 505 nm with a fluorescent microplate reader (Biotek, Winooski, VT, USA). Results were expressed as fold-change compared to control cells.

### Lactate dehydrogenase (LDH) assay

To perform the assay, 50μL of CyQUANT LDH Cytotoxicity Assay Kit reaction mixture was added to cell supernatants. After 30 minutes of incubation at room temperature, protected from light, the assay was stopped with a stop solution. Absorbance was measured at 490 nm and 680 nm using a microplate reader.

### Senescence associated β-galactosidase detection

To assess β-gal levels, a cellular senescence assay was performed (Dojindo, Kumamoto, Japan). After 24h of treatment with nintedanib, pirfenidone, or D +Q, cells were lysed with 50 μL of lysis buffer and incubated for 10 minutes. Then, 50 μL of SPiDER-βgal working solution was added to each well and incubated at 37 °C for 30 minutes

### Quantitative Real-Time Polymerase Chain Reaction (qRT-PCR)

Cells were lysed in Trizol^™^ reagent (Thermo-Fisher Scientific), and RNA was extracted as recommended by the manufacturer. 3μg of RNA was reverse transcribed into Complementary DNA (cDNA) using SuperScript^™^ II Reverse Transcriptase (Thermo-Fisher Scientific) as previously described (HOHMANN; HABIEL; ESPINDOLA; HUANG *et al*., 2021). Gene expression analyses were performed using TaqMan master mix (Thermo-Fisher Scientific) probes for human *Smooth Muscle Actin Alpha 2 (ACTA2), C-C Motif Chemokine Receptor (CCR)10, CDKN1A, CDKN2A, Collagen (COL)1A, COL3A1, (EPH Receptor A3) EPHA3, Fibronectin (FN)1, GDF15, and WNT16 (*all *Thermo-Fisher Scientific*). Quantitative PCR analysis was performed using Viia7 Thermocycler (Thermo-Fisher Scientific). Results were normalized to *RNA18S5* expression and presented as fold change values using DataAssist software version 3.01 (Thermo-Fisher Scientific).

### ELISA

IL-6, IL-8, Monocyte Chemoattractant Protein (MCP)-1, GDF-15 were determined in senescent fibroblast supernatant, and WNT16 levels were determined in senescent fibroblast lysates using a standardized sandwich Enzyme-Linked Immunoassay (ELISA) technique (R&D Systems, Minneapolis, MN, USA), according to manufacturer’s protocol.

### Soluble Collagen-1 direct ELISA

Senescent lung fibroblasts were plated into a 96 well plate and treated with nintedanib, pirfenidone, or D+Q for 24h. After 24 hours, lung fibroblast conditioned supernatants were harvested, and collagen-1 was assessed as previously described (HABIEL; ESPINDOLA; JONES; COELHO *et al*., 2018).

### Western blotting

Cells were lysed using RIPA lysis buffer (Thermo-Fisher Scientific) supplemented with Halt protease and phosphatase inhibitor cocktail (Thermo-Fisher Scientific). Protein concentrations were measured by using a Detergent Compatible protein assay (Bio-Rad Laboratories, Inc., Hercules, CA, USA), and the same amount of protein was loaded into a 4–15% NuPAGE Bis-Tris Protein gel. Gels were transferred using an iBlot Dry blotting system onto nitrocellulose membranes (Thermo-Fisher Scientific), and the transferred samples were blocked for 1 hour at room temperature in 5% non-fat-dry-milk in tris-buffered saline (TBS). Primary antibodies used included: Phospho-MLKL (Cat# 916895, Cell Signaling, Danvers, MA), Mixed Lineage Kinase Domain Like Pseudokinase (MLKL) (Cat #14993S, Cell Signaling), Gasdermin D (CAT#93709S, Cell Signaling), cleaved Gasdermin D (Cell Signaling CAT#36425S), and B cell lymphoma (Bcl)-2 (Cat#Ab182858, Abcam, Cambridge, UK). Images of chemiluminescent bands were acquired using a Bio-Rad Gel documentation system (Bio-Rad Laboratories, Inc.). Membranes were washed in TBS-T, blotted with anti-tubulin antibody (Abcam CAT#Ab6046), and developed similarly. Image Lab Software version 6.0.1 (Bio-Rad Laboratories, Inc.) was used to perform densitometric analysis.

### Data analysis

Statistical analyses were performed using GraphPad Prism 9.0.1 (GraphPad Software, San Diego, CA, USA). Data were expressed as mean ± Standard Error of the Mean (SEM) and evaluated for significance by one-way Analysis of Variance (ANOVA) followed by Tukey’s test. A value of P < 0.05 was considered statistically significant.

## Results

### SOC drugs do not alter cell viability, LDH leakage, or caspase-3 release

To explore the hypothesis that SOC drugs modulate apoptosis in senescent normal and IPF fibroblasts, we treated these cells with nintedanib (300nM), pirfenidone (2.5mM), D+Q (20μM /15μM), or vehicle (Dimethylsulfoxide (DMSO) 0.05%) for 24h, followed by the addition of Super FasL (100 ng/mL) or TRAIL (100ng/ml) + His-tag (5μg/ml). Neither nintedanib nor pirfenidone showed an effect on cell viability or LDH leakage, on normal or IPF senescent lung fibroblasts. However, nintedanib increased caspase-3 activity when combined with the cell death ligand FasL only on normal cells (Figure 1C).

**Figure 1.**
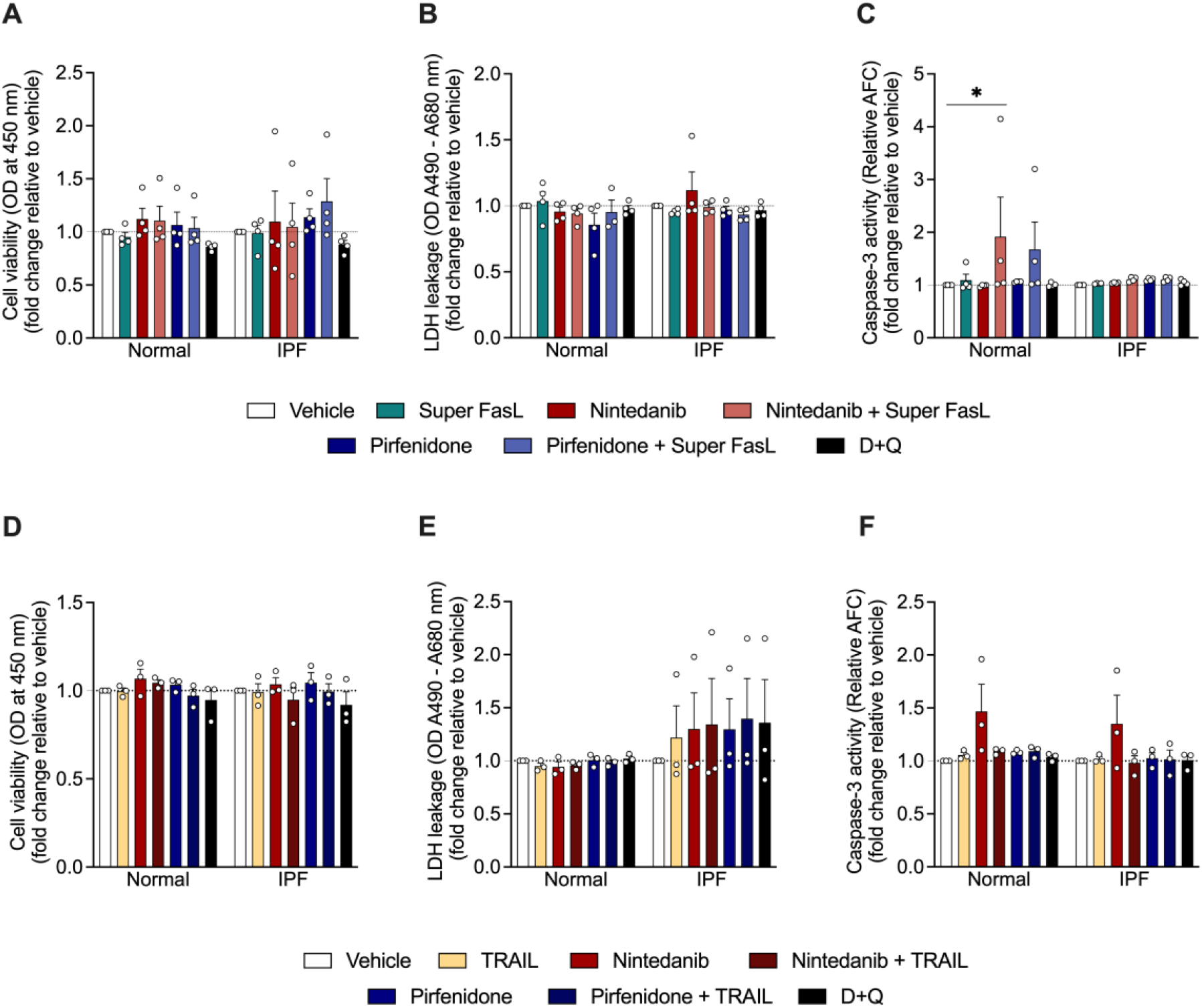
Cell viability, lactate dehydrogenase (LDH) release, and caspase-3 activity by senescent lung fibroblasts from normal and IPF patients after 24-hour treatment with nintedanib (300 nM), pirfenidone (2.5 mM), Dasatinib (20 μM) + Quercetin (15 μM), or vehicle (DMSO 0.05%), followed by 3h of Super Fas Ligand (Super FasL, 100 ng/ml) or Tumor necrosis factor (TNF)-related apoptosis-inducing ligand (TRAIL, 100 ng/ml) + His-tag (5 μg/ml). Cell viability was expressed as optical density (OD) values at 450 nm, LDH release as the difference in the absorbance values at 490 and 680 nm, and caspase-3 activity as relative 7-amino-4-trifluoromethyl coumarin (AFC). Data are presented as mean ± SEM (n = 3 or 4 per group). *p<0.05.

### Expression of marker of senescence in normal and IPF senescent fibroblasts

Normal and IPF lung fibroblasts were submitted to serial passage rounds until cells obtained the senescence phenotype, as previously described (HOHMANN; HABIEL; COELHO; VERRI *et al*., 2019). One of the most used methods to evaluate cellular senescence is the detection of β-GAL activity (DE MERA-RODRIGUEZ; ALVAREZ-HERNAN; GANAN; MARTIN-PARTIDO *et al*., 2021). We confirmed that cells are beta-galactosidase positive by measuring Spider-βGal intensity, however, we did not find any significant difference after the treatment with SOC drugs or D+Q (Figure 2A). We next determined the expression of *CDKN1A* and *CDKN2A* used as markers of cellular senescence (TOMINAGA, 2015). We observed a reduction of *CDKN1A* mRNA expression only in normal lung fibroblasts after the treatment with nintedanib (Figure 2B-C). In addition, GDF15 has been found as one of the components of SASP (ZHANG; JIANG; NOURAIE; ROTH *et al*., 2019). We observed that *GDF15* was upregulated after D+Q treatment in IPF fibroblasts, which was not observed in any of the other treatments (Figure 2D). Moreover, Wnt16 emerged as a new marker of senescence, regulating p53 activity and phosphatidylinositol 3-kinase (PI3K)/Ak Strain Transforming (AKT) pathway (BINET; YTHIER; ROBLES; COLLADO *et al*., 2009). We did not observe any significant difference in *WNT16* mRNA expression among the treated groups (Figure 2E).

**Figure 2.**
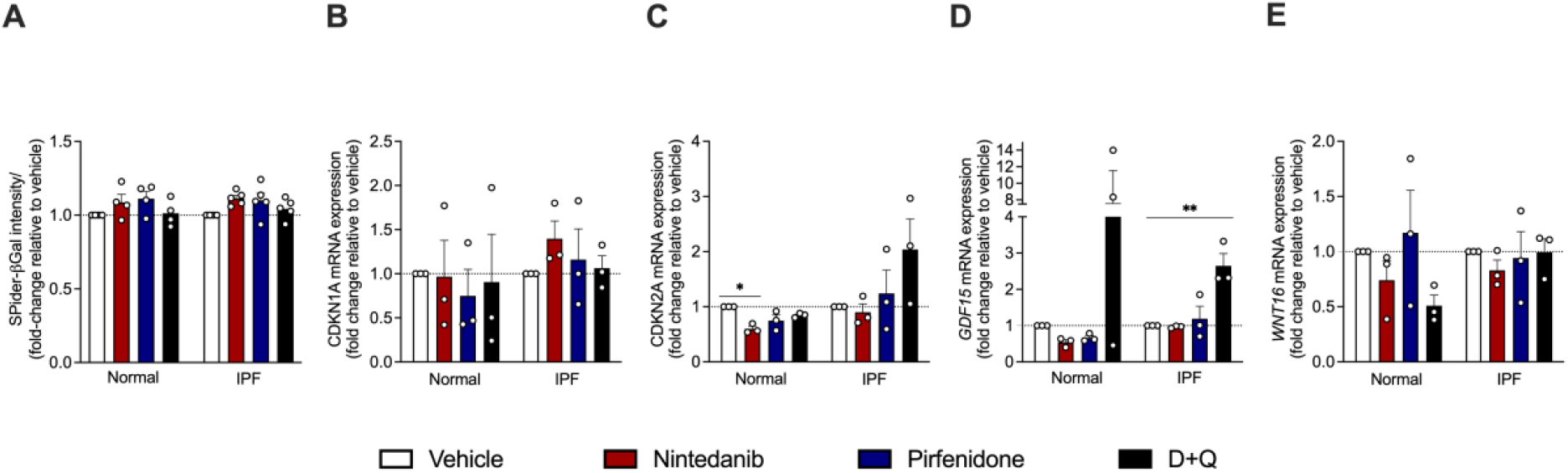
Lung fibroblasts isolated from normal and patients with idiopathic pulmonary fibrosis (IPF) exhibited a senescent phenotype. Fibroblasts isolated from the lungs of normal patients or IPF patients were plated simultaneously at time 0 and treated with nintedanib (300 nM), pirfenidone (2.5 mM), and D+Q (20 μM/15 μM) or vehicle (DMSO 0.05%) for 24h. *CDKN1A, CDKN2A, GDF15*, and *WNT16* transcripts were first normalized to the housekeeping gene *18S*. Data are presented as mean ± SEM (n = 3-5 per group). *P<0.05, **P<0.001.

### Influence of SOC and D+Q on senescent lung fibroblasts

We next examined whether SOC drugs and D+Q had an effect on the expression of *ACTA2, CCR10, EPHA3, COL3A*, and *FN1*. The treatment with nintedanib was able to reduce *ACTA2* mRNA expression only in normal senescent lung fibroblast, while the other drugs did not show an effect on *ACTA2* expression (Figure 3A). Moreover, all drugs reduced *COL3A1* mRNA expression in normal senescent lung fibroblast. However, the same effect was not observed in IPF cells. The deposition of extracellular matrix proteins, like collagen and fibronectin in the lung, triggers respiratory failure (BUENO; CALYECA; ROJAS; MORA, 2020). The treatment with pirfenidone enhanced *FN1* in IPF senescent lung fibroblast. Nintedanib and D+Q reduced *EPHA3*, a mesenchymal cell marker in normal senescent lung fibroblast (Figure 3)

**Figure 3.**
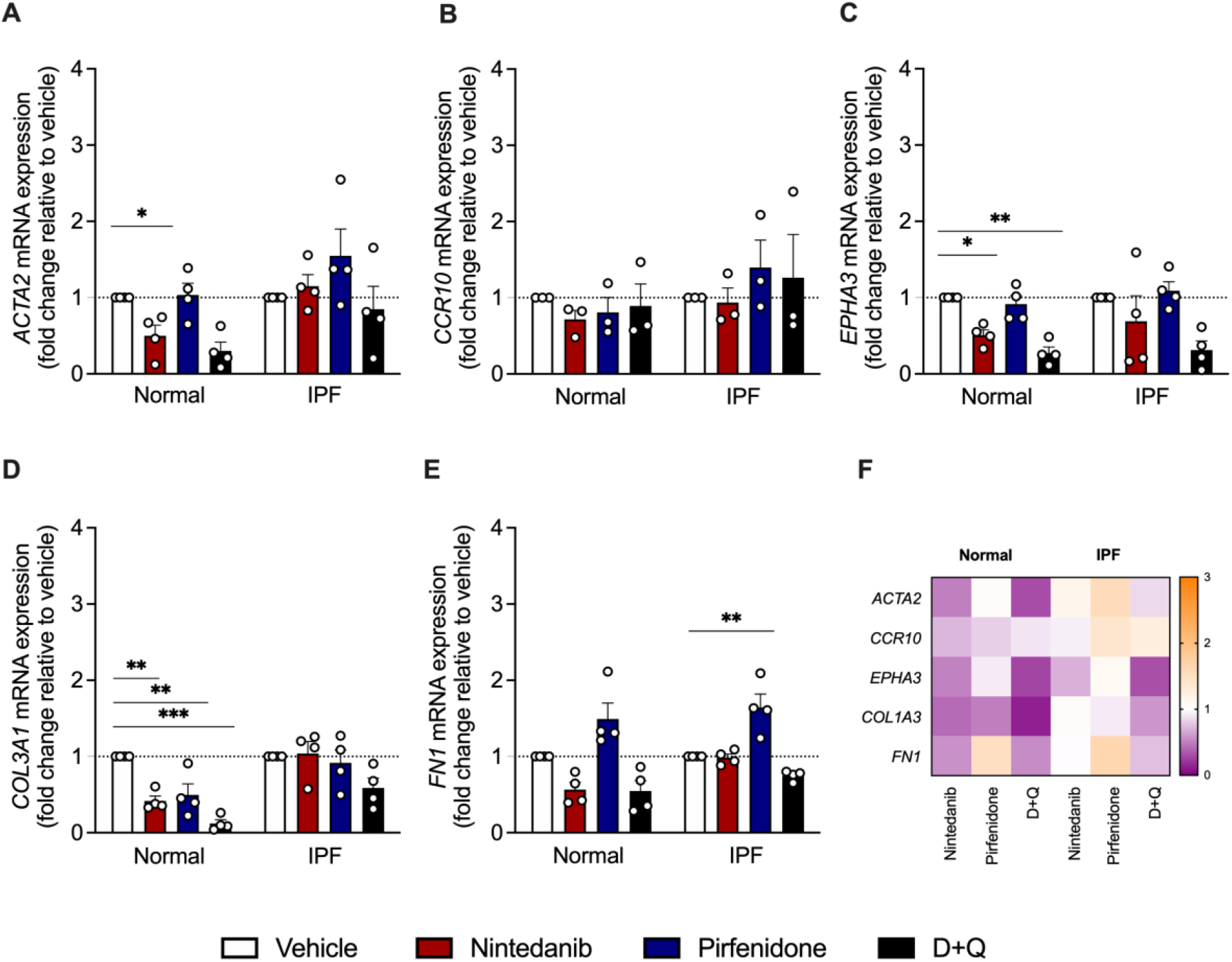
Effects of SOC drugs and D+Q on *ACTA2, CCR10, EPHA3, COL3A1*, and *FN1* mRNA expression in normal and IPF senescent lung fibroblasts. Heatmap of the expression of SASP and fibrosis-related genes in lung fibroblasts from normal and IPF patients treated with nintedanib (300 nM), pirfenidone (2.5 mM), D+Q (20 μM/15 μM) for 24h. Each transcript was first normalized to the housekeeping gene RNA *18S*. Upregulation (orange) and downregulation (purple) of gene expression, compared with vehicle-treated cells. Data are presented as mean ± SEM (n= 3 or 4 per group). *P<0.05; **P<0.001; and ***P<0.0001.

### Pirfenidone increases COL1A1 expression in senescent IPF fibroblasts, and D+Q reduces collagen expression and release in senescent normal and IPF fibroblasts

One of the fibrosis hallmarks is the excessive deposition of fibrotic extracellular matrix proteins, especially type I collagen (KLEAVELAND; VELIKOFF; YANG; AGARWAL *et al*., 2014). To determine the effects of SOC drugs on the fibrosis-related marker type I collagen, we investigated the expression of *COL1A1* in senescent lung fibroblasts after the treatment with nintedanib, pirfenidone, or D+Q. RT-PCR and ELISA demonstrated that the expression levels of *COL1A1* were significantly increased in the IPF group after pirfenidone treatment (Figure 4A). On the other hand, D+Q treatment significantly decreased type I collagen secretion in normal and IPF lung fibroblasts (Figure 4B).

**Figure 4.**
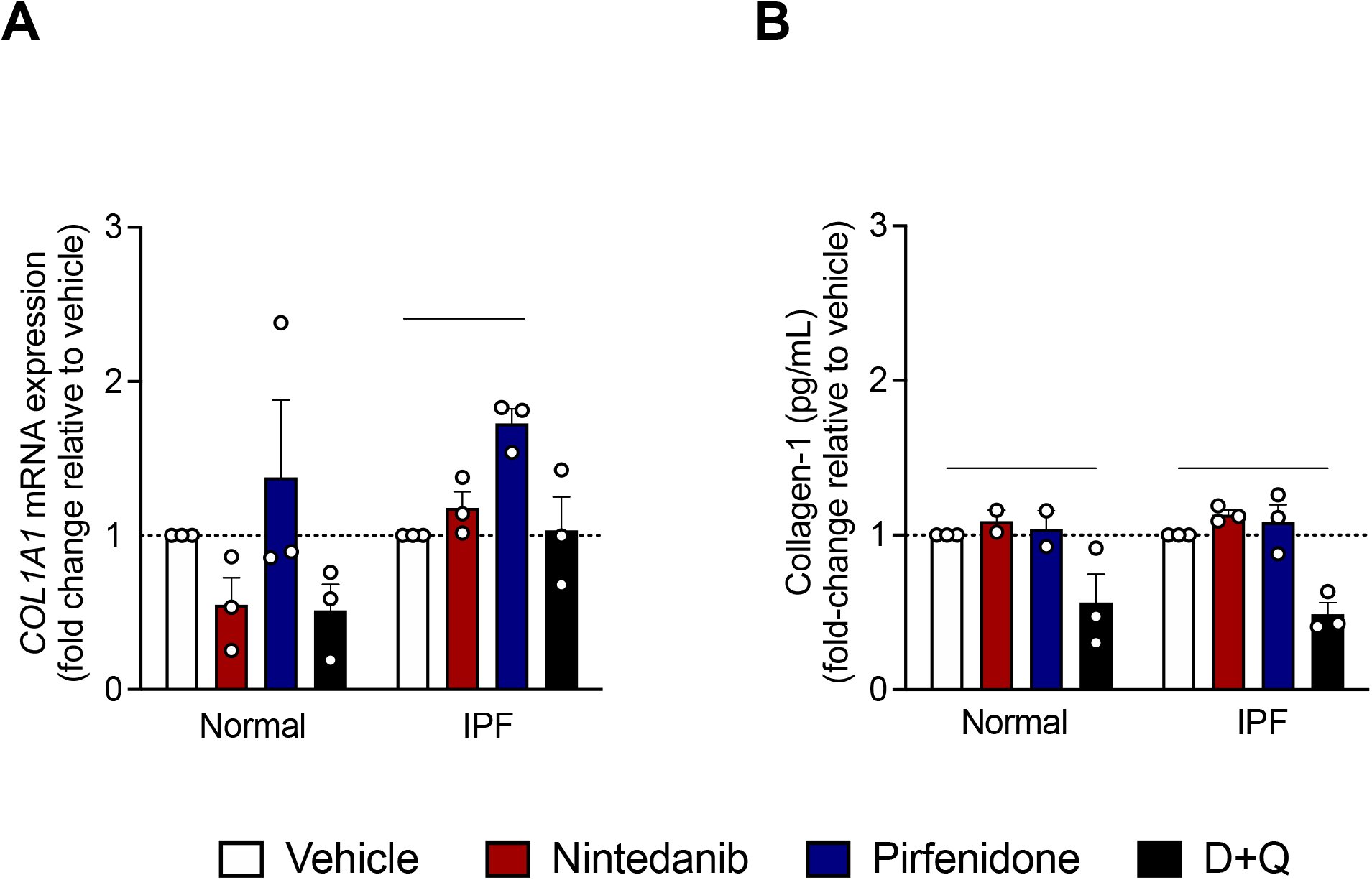
Influence of SOC drugs on COL1A1 mRNA expression and type I collagen secretion. Effects of nintedanib (300nM), pirfenidone (2.5mM), D+Q (20μM/15μM) or vehicle (DMSO 0.05%), on *COL1A1* mRNA expression and collagen-1 levels. Data represented as mean ± SEM (n = 3 per group), *p<0.05. **p<0.01.

**Figure 4.**
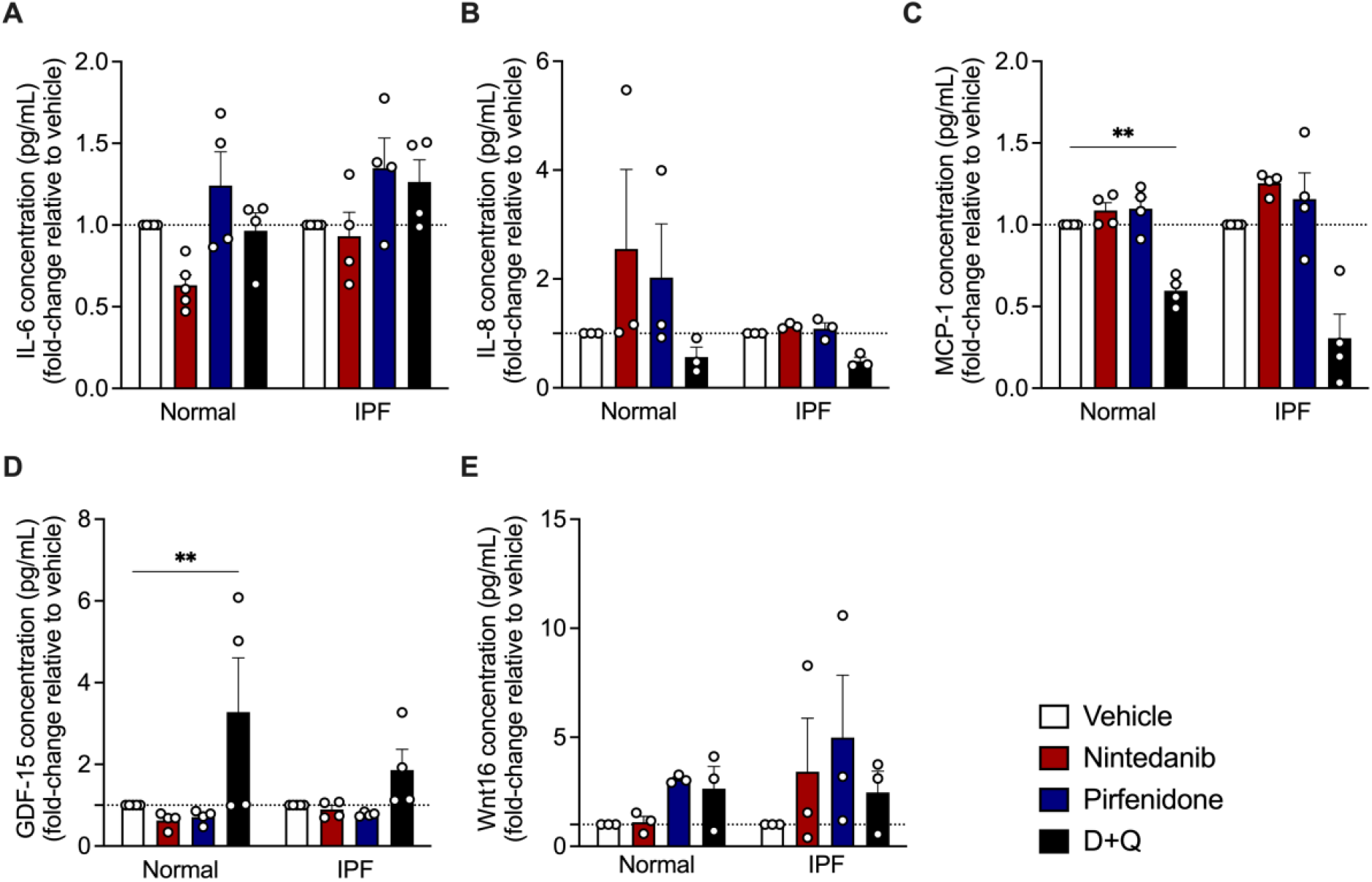
SOC do not have an impact on SASP release. Effects of nintedanib (300nM), pirfenidone (2.5mM), dasatinib (20μM) + quercetin (15μM), or control (vehicle; DMSO 0.05%), on IL-6, IL-8, MCP-1, GDF-15, and Wnt16 levels. Data are presented as mean ± SEM (n= 3 or 4 per group). **P<0.001

### SOC drugs influence IL-6, IL-8, MCP-1, GDF-15, Wnt16 in senescent cells

Next, we investigated whether SOC drugs affect senescent cells via modulation of their secretome. Senescent cells secrete interleukins, inflammatory cytokines, and growth factors that can affect neighboring cells (COPPE; DESPREZ; KRTOLICA; CAMPISI, 2010). Among SASP cytokines, the pleiotropic pro-inflammatory cytokine IL-6 is the most distinguished. We observed that SOC drugs do not affect IL-6 and IL-8 in senescent normal and IPF lung fibroblasts. Taking into consideration that senescent fibroblast can recruit leukocytes, and MCP-1 is a crucial component of SASP (JIN; LEE; HEO; LIM *et al*., 2016), we evaluated the concentration of MCP-1 on normal and IPF senescent fibroblasts. The treatment with D+Q was able to reduce MCP-1 release in normal but not in IPF fibroblast, while SOC drugs showed no effect neither in normal nor IPF senescent fibroblast. In addition, compelling studies showed GDF-15 emerging as part of the SASP repertoire (AL-MUDARES; REDDICK; REN; VENKATESH *et al*., 2020). After the dosage of GDF-15, we observed a prominent increase after D+Q treatment only in normal senescent fibroblasts. However, no significant results were observed after the treatment with SOC drugs in normal or IPF senescent cells. Moreover, we observed that after the treatment with SOC drugs and D+Q, there was no significant difference in Wnt16 release when compared with vehicle, showing that those drugs do not affect Wnt16 (Figure 5).

**Figure 5.**
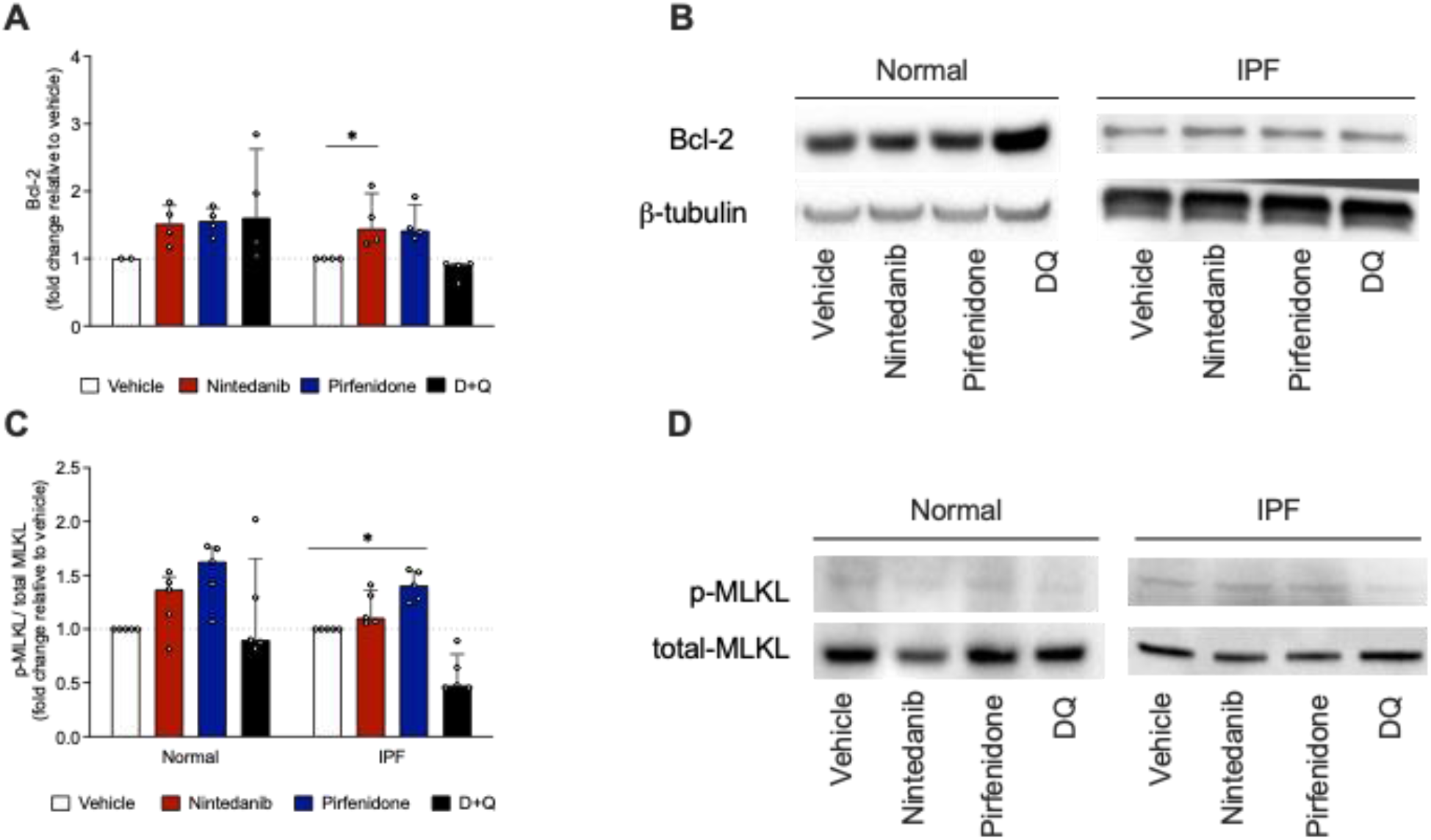
Effects of SOC drugs on Bcl-2, phosphorylation of MLKL, and cathepsin B release. (A) Western blot quantification of Bcl-2. (B) Representative western blot of Bcl-2. (C) Western blot quantification of MLKL phosphorylated/MLKL total (D) Representative western blot of p-MLKL/MLKL protein expression. Data are presented as median with interquartile range, one-way ANOVA followed by Tukey’s test. *p<0.05 **p<0.01 and p<0.001.

### SOC drugs trigger necroptosis but not pyroptosis

Because previous experiments have shown that SOC drugs do not trigger apoptosis, we next evaluated the influence of SOC drugs on the apoptosis regulator Bcl-2. We saw an increase in Bcl-2 protein levels after the treatment with nintedanib (Figure 5A), and the same effect was not observed in cells treated with pirfenidone or D+Q. Subsequently, to address whether SOC drugs trigger other cell death types, we performed western blot to measure phosphorylated MLKL (p-MLKL) and total MLKL in senescent fibroblasts from normal and IPF patients. The activation of MLKL upon its phosphorylation initiates necroptosis, a form of programmed cell death in which rupture of cellular membranes leads to the release of intracellular components (YOON; KOVALENKO; BOGDANOV; WALLACH, 2017). Treatment with pirfenidone significantly increases the ratio p-MLKL/MLKL (Figure 6C-D), concluding that this drug leads to necroptosis. Although Nintedanib presented a mild increase of p-MLKL/MLKL, it was not significant when compared to the vehicle. Another cell death type evaluated was pyroptosis, which is triggered by pro-inflammatory signals and associated with inflammation (BERTHELOOT; LATZ; FRANKLIN, 2021). We evaluated the presence of cleaved Gasdermin D. However, it was not detected either in senescent fibroblast after the treatment with vehicle, SOC drugs, or D+Q, confirming that such drugs do not lead to pyroptosis (data not shown).

**Figure 6.**
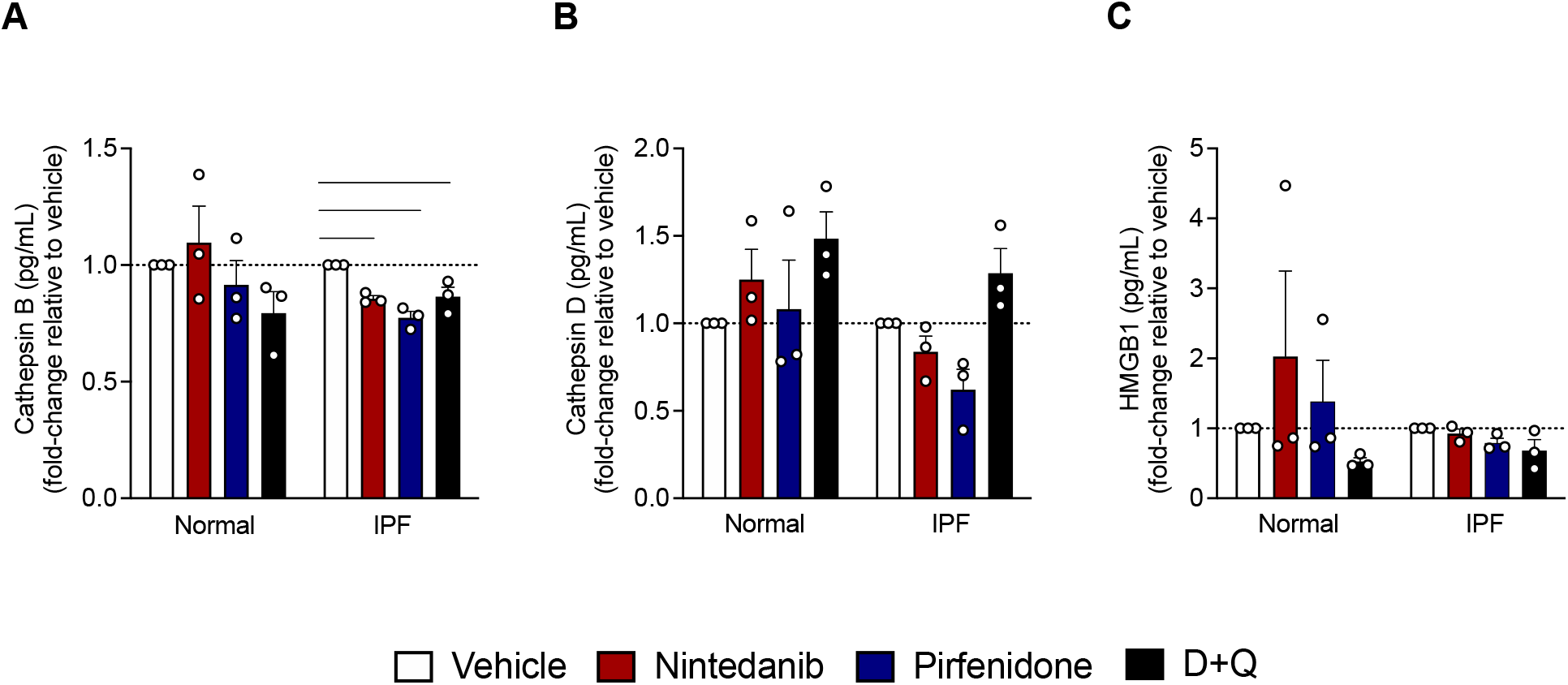
Necroptosis secreted factors after treatment with SOC drugs. Effects of Nintedanib (300nM), Pirfenidone (2.5mM), Dasatinib (20μM) + Quercetin (15μM) or control (vehicle; DMSO 0.05%), on cathepsin B, cathepsin D and (HMGB1) levels. Data are presented as mean ± SEM (n= 3 per group). *P<0.05 and ***P<0.0001 as indicated by the bars.

### SOC drugs alter cathepsin B release but not cathepsin D or High Mobility Group Box 1 (HMGB1)

We next evaluated the role of SOC drugs in the release of cathepsin B, cathepsin D, and HMGB1 release. We observed that the treatment with nintedanib and pirfenidone was able to reduce the levels of cathepsin B in IPF senescent lung fibroblast, while no significant difference was observed in the normal group. Moreover, any of the treatments influenced cathepsin D and HMGB1 release (Figure 6).

## Discussion

Senescence is a well-established feature of IPF (MEINERS; LEHMANN, 2020). Senescent cells implicate in the generation of age-related phenotypes, and their removal can prevent or delay tissue dysfunction and lengthen healthspan (BAKER; WIJSHAKE; TCHKONIA; LEBRASSEUR *et al*., 2011). In addition, the clearance of senescence cells has been shown to protect mice from lung fibrosis (LEHMANN; KORFEI; MUTZE; KLEE *et al*., 2017). Under this senescent state, the cells secrete a myriad of cytokines and proteases and growth factors that play a pivotal impact on adjacent cells and the tissue microenvironment (KIRKLAND; TCHKONIA, 2017). Notwithstanding, senescent cells are remarkably resistant to apoptosis due to the upregulation of proteins of anti-apoptotic pathways (HU; LI; ZI; LI *et al*., 2022).

Mounting evidence has shown that the progression of fibrosis involves ECM-driven mechanisms. Mesenchymal cells and their ECM products lead to the expansion of the alveolar wall, causing loss of the gas-exchange surface. The myofibroblast core of IPF presents components such as type I and III collagens, extra domain fibronectin, and fibrin that are related to the progression of IPF (HERRERA; HENKE; BITTERMAN, 2018). We observed an increase of α-SMA, collagen-1, and fibronectin 1 after the treatment with pirfenidone, suggesting that although evidence shows its anti-fibrotic effect in proliferative fibroblasts (COLLINS; RAGHU, 2019), the same effect is not perceived in senescent fibroblasts. In agreement with our findings, Roach et al. (2021) and collaborators found an increased expression of α-SMA, deposition of collagen 1 and collagen-3, and secretion of soluble collagen after treating fibroblasts with nintedanib and pirfenidone (ROACH; CASTELLS; DIXON; MASON *et al*., 2021).

In a bleomycin model, GDF-15 showed to be the most upregulated protein and suggested as a suitable biomarker of epithelial stress and severity of the disease (ZHANG; JIANG; NOURAIE; ROTH *et al*., 2019). Moreover, after a proteomics analysis Tanaka et al. found a positive association between GDF15 with age (AL-MUDARES; REDDICK; REN; VENKATESH *et al*., 2020). In addition, GDF15 is reported to increase α-SMA expression in WI-38 lung fibroblasts, evoking the hypothesis that elevated GDF15 in the fibrotic lung is involved in fibroblast activation (TAKENOUCHI; KITAKAZE; TSUBOI; OKAMOTO, 2020). After D+Q treatment, GDF-15 levels were significantly higher when compared to SOC drugs, suggesting that D+Q could be triggering the stage-specific embryonic antigen (SSEA)-4+ cell to proliferate and trigger the emersion of resistant cells.

In the present study, we showed that the treatment with SOC drugs did not evoke apoptosis in senescent fibroblasts either spontaneously or via ligands induced. The resistance to apoptosis has been demonstrated by Moodley et al., they showed *in vitro* that myofibroblasts isolated from fibrotic lungs were more resistant to Fas ligand-induced apoptosis than controls (MOODLEY; CATERINA; SCAFFIDI; MISSO *et al*., 2004). Moreover, the accumulation of proteins from the Bcl-2 family contributes to the resistance to undergoing apoptosis (YOSEF; PILPEL; TOKARSKY-AMIEL; BIRAN *et al*., 2016). We observed that senescent IPF cells treated with nintedanib presented elevated levels of Bcl-2 when compared to normal senescent cells, despite previous studies showing nintedanib inhibits apoptotic proteins in lung-resident myofibroblasts (KASAM; REDDY; JEGGA; MADALA, 2019), although the effect was not evaluated in senescent cells, which are naturally resistant to apoptosis (SALMINEN; OJALA; KAARNIRANTA, 2011).

It has been demonstrated augmented levels of Receptor Interacting Serine/Threonine Kinase 3 (RIPK3) and phosphorylated MLKL in alveolar epithelial cells (AECs) in IPF lungs (LEE; YOSHIDA; KIM; LEE *et al*., 2018). Lee et al. postulated that the involvement of necroptosis in IPF pathogenesis might be in response to cell-damaging agents’ injury to AECs, leading to apoptosis and RIPK-3–regulated necroptosis. Damage-associated molecular patterns, including HMGB1 and IL-β, released from necroptotic AECs are responsible not only for inflammation but also for fibrosis development through enhanced myofibroblast differentiation during IPF pathogenesis.

Taken together that caspase-3 release was not detected, we investigated whether other kinds of cell death could be involved. Necroptosis is an alternative system for cell death when caspase-dependent apoptosis is restricted or absent (HAN; ZHONG; ZHANG, 2011). It is present in many pathologies such as inflammatory diseases (TAO; SUN; WU; WANG *et al*., 2020), ischemia-reperfusion injuries (MULLER; DEWITZ; SCHMITZ; SCHRODER *et al*., 2017), and degenerative diseases (GAUTHERON; VUCUR; SCHNEIDER; SEVERI *et al*., 2016). The necroptosis pathway can be induced by impaired apoptosis by ligand-dependent stimulation of cell death receptors, for instance, Fas (CHOI; PRICE; RYTER; CHOI, 2019). It is orchestrated by distinct proteins, namely RIPK1 and RIPK3 and the downstream protein MLKL, that once phosphorylated by RIPK3, lead to necroptosis by inducing the formation of oligomers, migration to the plasma membrane, and binding to phosphatidylinositol lipids to directly disrupt membrane integrity (RODRIGUEZ; WEINLICH; BROWN; GUY *et al*., 2016). In the present study, we found some evidence that pirfenidone increases MLKL phosphorylation, the major hallmark of necroptosis, and the same effect was not observed after the treatment with nintedanib or D+Q. However, the mechanism whereby pirfenidone enhances this phosphorylation requires further investigation.

Previous work has shown that the inhibition of cathepsins provokes the induction of necroptosis in bone marrow-derived macrophages, suggesting that cathepsins act as anti-necroptotic factors (MCCOMB; SHUTINOSKI; THURSTON; CESSFORD *et al*., 2014). This finding corroborates with our study, we found a decrease in cathepsin B after the treatment with nintedanib, pirfenidone, and, interestingly, D+Q. Similar results were obtained with cathepsin D. Moreover, cathepsins are involved in the degradation of the anti-apoptotic protein Bcl-2, leading to apoptosis (DROGA-MAZOVEC; BOJIC; PETELIN; IVANOVA *et al*., 2008). However, SOC drugs showed to diminish cathepsin activity, which triggers an evasion of apoptosis and the increase of necroptosis by the phosphorylation of MLKL.

Together, these data demonstrate that with IPF progression and the accumulation of senescence cells, SOC drugs show insufficient effects on these cells, and are not able to reduce fibrotic markers. Moreover, Pirfenidone showed to promote a inflamatory cell death, however the mechanism which this drug mediates necroptosis remains elusive. Overall, this study sheds light on the need to seek more effective therapies that can target and selectively eliminate senescence cells in IPF, and further studies are still necessary to reveal SOC drugs’ effects and mechanism of action on senescent cells.

## Acknowledgements

WAV reports CNPq fellowship (#309633/2021-4).

## References

1. Martinez FJ, Collard HR, Pardo A, Raghu G, Richeldi L, Selman M, Swigris JJ, Taniguchi H, Wells AU. Idiopathic pulmonary fibrosis. Nat Rev Dis Primers 2017; 3: 17074.

2. Raghu G, Chen SY, Hou Q, Yeh WS, Collard HR. Incidence and prevalence of idiopathic pulmonary fibrosis in US adults 18-64 years old. Eur Respir J 2016; 48: 179–186.

3. King TE Jr.,, Pardo A, Selman M. Idiopathic pulmonary fibrosis. Lancet 2011; 378: 1949–1961.

4. Maher TM, Kreuter M, Lederer DJ, Brown KK, Wuyts W, Verbruggen N, Stutvoet S, Fieuw A, Ford P, Abi-Saab W, Wijsenbeek M. Rationale, design and objectives of two-phase III, randomised, placebo-controlled studies of GLPG1690, a novel autotaxin inhibitor, in idiopathic pulmonary fibrosis (ISABELA 1 and 2). BMJ Open Respir Res 2019; 6: e000422.

5. American Thoracic Society. Idiopathic pulmonary fibrosis: diagnosis and treatment. International consensus statement. American Thoracic Society (ATS) and the European Respiratory Society (ERS). Am J Respir Crit Care Med 2000; 161: 646–664.

6. Salama R, Sadaie M, Hoare M, Narita M. Cellular senescence and its effector programs. Genes Dev 2014; 28: 99–114.

7. Kendall RT, Feghali-Bostwick CA. Fibroblasts in fibrosis: novel roles and mediators. Front Pharmacol 2014; 5: 123.

8. Parimon T, Hohmann MS, Yao C. Cellular Senescence: Pathogenic Mechanisms in Lung Fibrosis. Int J Mol Sci 2021; 22.

9. Baker DJ, Childs BG, Durik M, Wijers ME, Sieben CJ, Zhong J, Saltness RA, Jeganathan KB, Verzosa GC, Pezeshki A, Khazaie K, Miller JD, van Deursen JM. Naturally occurring p16(Ink4a)-positive cells shorten healthy lifespan. Nature 2016; 530: 184–189.

10. Soto-Gamez A, Quax WJ, Demaria M. Regulation of Survival Networks in Senescent Cells: From Mechanisms to Interventions. J Mol Biol 2019; 431: 2629–2643.

11. Campisi J, d’Adda di Fagagna F. Cellular senescence: when bad things happen to good cells. Nat Rev Mol Cell Biol 2007; 8: 729–740.

12. Faget DV, Ren Q, Stewart SA. Unmasking senescence: context-dependent effects of SASP in cancer. Nat Rev Cancer 2019; 19: 439–453.

13. Binet R, Ythier D, Robles AI, Collado M, Larrieu D, Fonti C, Brambilla E, Brambilla C, Serrano M, Harris CC, Pedeux R. WNT16B is a new marker of cellular senescence that regulates p53 activity and the phosphoinositide 3-kinase/AKT pathway. Cancer Res 2009; 69: 9183–9191.

14. Basisty N, Kale A, Jeon OH, Kuehnemann C, Payne T, Rao C, Holtz A, Shah S, Sharma V, Ferrucci L, Campisi J, Schilling B. A proteomic atlas of senescence-associated secretomes for aging biomarker development. PLoS Biol 2020; 18: e3000599.

15. Karimi-Shah BA, Chowdhury BA. Forced vital capacity in idiopathic pulmonary fibrosis--FDA review of pirfenidone and nintedanib. N Engl J Med 2015; 372: 1189–1191.

16. Raghu G, Selman M. Nintedanib and pirfenidone. New antifibrotic treatments indicated for idiopathic pulmonary fibrosis offer hopes and raises questions. Am J Respir Crit Care Med 2015; 191: 252–254.

17. Hostettler KE, Zhong J, Papakonstantinou E, Karakiulakis G, Tamm M, Seidel P, Sun Q, Mandal J, Lardinois D, Lambers C, Roth M. Anti-fibrotic effects of nintedanib in lung fibroblasts derived from patients with idiopathic pulmonary fibrosis. Respir Res 2014; 15: 157.

18. Kato K, Shin YJ, Palumbo S, Papageorgiou I, Hahn S, Irish JD, Rounseville SP, Krafty RT, Wollin L, Sauler M, Hecker L. Leveraging ageing models of pulmonary fibrosis: the efficacy of nintedanib in ageing. Eur Respir J 2021; 58.

19. Kellogg DL, Kellogg DL Jr.,, Musi N, Nambiar AM. Cellular Senescence in Idiopathic Pulmonary Fibrosis. Curr Mol Biol Rep 2021; 7: 31–40.

20. Schafer MJ, White TA, Iijima K, Haak AJ, Ligresti G, Atkinson EJ, Oberg AL, Birch J, Salmonowicz H, Zhu Y, Mazula DL, Brooks RW, Fuhrmann-Stroissnigg H, Pirtskhalava T, Prakash YS, Tchkonia T, Robbins PD, Aubry MC, Passos JF, Kirkland JL, Tschumperlin DJ, Kita H, LeBrasseur NK. Cellular senescence mediates fibrotic pulmonary disease. Nat Commun 2017; 8: 14532.

21. Justice JN, Nambiar AM, Tchkonia T, LeBrasseur NK, Pascual R, Hashmi SK, Prata L, Masternak MM, Kritchevsky SB, Musi N, Kirkland JL. Senolytics in idiopathic pulmonary fibrosis: Results from a first-in-human, open-label, pilot study. EBioMedicine 2019; 40: 554–563.

22. Hohmann MS, Habiel DM, Coelho AL, Verri WA Jr.,, Hogaboam CM. Quercetin Enhances Ligand-induced Apoptosis in Senescent Idiopathic Pulmonary Fibrosis Fibroblasts and Reduces Lung Fibrosis In Vivo. Am J Respir Cell Mol Biol 2019; 60: 28–40.

23. Hohmann MS, Habiel DM, Espindola MS, Huang G, Jones I, Narayanan R, Coelho AL, Oldham JM, Noth I, Ma SF, Kurkciyan A, McQualter JL, Carraro G, Stripp B, Chen P, Jiang D, Noble PW, Parks W, Woronicz J, Yarranton G, Murray LA, Hogaboam CM. Antibody-mediated depletion of CCR10+EphA3+ cells ameliorates fibrosis in IPF. JCI Insight 2021; 6.

24. Habiel DM, Espindola MS, Jones IC, Coelho AL, Stripp B, Hogaboam CM. CCR10+ epithelial cells from idiopathic pulmonary fibrosis lungs drive remodeling. JCI Insight 2018; 3.

25. de Mera-Rodriguez JA, Alvarez-Hernan G, Ganan Y, Martin-Partido G, Rodriguez-Leon J, Francisco-Morcillo J. Is Senescence-Associated beta-Galactosidase a Reliable in vivo Marker of Cellular Senescence During Embryonic Development? Front Cell Dev Biol 2021; 9: 623175.

26. Tominaga K. The emerging role of senescent cells in tissue homeostasis and pathophysiology. Pathobiol Aging Age Relat Dis 2015; 5: 27743.

27. Bueno M, Calyeca J, Rojas M, Mora AL. Mitochondria dysfunction and metabolic reprogramming as drivers of idiopathic pulmonary fibrosis. Redox Biol 2020; 33: 101509.

28. Kleaveland KR, Velikoff M, Yang J, Agarwal M, Rippe RA, Moore BB, Kim KK. Fibrocytes are not an essential source of type I collagen during lung fibrosis. J Immunol 2014; 193: 5229–5239.

29. Coppe JP, Desprez PY, Krtolica A, Campisi J. The senescence-associated secretory phenotype: the dark side of tumor suppression. Annu Rev Pathol 2010; 5: 99–118.

30. Jin HJ, Lee HJ, Heo J, Lim J, Kim M, Kim MK, Nam HY, Hong GH, Cho YS, Choi SJ, Kim IG, Shin DM, Kim SW. Senescence-Associated MCP-1 Secretion Is Dependent on a Decline in BMI1 in Human Mesenchymal Stromal Cells. Antioxid Redox Signal 2016; 24: 471–485.

31. Al-Mudares F, Reddick S, Ren J, Venkatesh A, Zhao C, Lingappan K. Role of Growth Differentiation Factor 15 in Lung Disease and Senescence: Potential Role Across the Lifespan. Front Med (Lausanne) 2020; 7: 594137.

32. Yoon S, Kovalenko A, Bogdanov K, Wallach D. MLKL, the Protein that Mediates Necroptosis, Also Regulates Endosomal Trafficking and Extracellular Vesicle Generation. Immunity 2017; 47: 51–65 e57.

33. Meiners S, Lehmann M. Senescent Cells in IPF: Locked in Repair? Front Med (Lausanne) 2020; 7: 606330.

34. Baker DJ, Wijshake T, Tchkonia T, LeBrasseur NK, Childs BG, van de Sluis B, Kirkland JL, van Deursen JM. Clearance of p16Ink4a-positive senescent cells delays ageing-associated disorders. Nature 2011; 479: 232–236.

35. Lehmann M, Korfei M, Mutze K, Klee S, Skronska-Wasek W, Alsafadi HN, Ota C, Costa R, Schiller HB, Lindner M, Wagner DE, Gunther A, Konigshoff M. Senolytic drugs target alveolar epithelial cell function and attenuate experimental lung fibrosis ex vivo. Eur Respir J 2017; 50.

36. Kirkland JL, Tchkonia T. Cellular Senescence: A Translational Perspective. EBioMedicine 2017; 21: 21–28.

37. Hu L, Li H, Zi M, Li W, Liu J, Yang Y, Zhou D, Kong QP, Zhang Y, He Y. Why Senescent Cells Are Resistant to Apoptosis: An Insight for Senolytic Development. Front Cell Dev Biol 2022; 10: 822816.

38. Herrera J, Henke CA, Bitterman PB. Extracellular matrix as a driver of progressive fibrosis. J Clin Invest 2018; 128: 45–53.

39. Collins BF, Raghu G. Antifibrotic therapy for fibrotic lung disease beyond idiopathic pulmonary fibrosis. Eur Respir Rev 2019; 28.

40. Roach KM, Castells E, Dixon K, Mason S, Elliott G, Marshall H, Poblocka MA, Macip S, Richardson M, Khalfaoui L, Bradding P. Evaluation of Pirfenidone and Nintedanib in a Human Lung Model of Fibrogenesis. Front Pharmacol 2021; 12: 679388.

41. Zhang Y, Jiang M, Nouraie M, Roth MG, Tabib T, Winters S, Chen X, Sembrat J, Chu Y, Cardenes N, Tuder RM, Herzog EL, Ryu C, Rojas M, Lafyatis R, Gibson KF, McDyer JF, Kass DJ, Alder JK. GDF15 is an epithelial-derived biomarker of idiopathic pulmonary fibrosis. Am J Physiol Lung Cell Mol Physiol 2019; 317: L510–L521.

42. Takenouchi Y, Kitakaze K, Tsuboi K, Okamoto Y. Growth differentiation factor 15 facilitates lung fibrosis by activating macrophages and fibroblasts. Exp Cell Res 2020; 391: 112010.

43. Moodley YP, Caterina P, Scaffidi AK, Misso NL, Papadimitriou JM, McAnulty RJ, Laurent GJ, Thompson PJ, Knight DA. Comparison of the morphological and biochemical changes in normal human lung fibroblasts and fibroblasts derived from lungs of patients with idiopathic pulmonary fibrosis during FasL-induced apoptosis. J Pathol 2004; 202: 486–495.

44. Yosef R, Pilpel N, Tokarsky-Amiel R, Biran A, Ovadya Y, Cohen S, Vadai E, Dassa L, Shahar E, Condiotti R, Ben-Porath I, Krizhanovsky V. Directed elimination of senescent cells by inhibition of BCL-W and BCL-XL. Nat Commun 2016; 7: 11190.

45. Kasam RK, Reddy GB, Jegga AG, Madala SK. Dysregulation of Mesenchymal Cell Survival Pathways in Severe Fibrotic Lung Disease: The Effect of Nintedanib Therapy. Front Pharmacol 2019; 10: 532.

46. Salminen A, Ojala J, Kaarniranta K. Apoptosis and aging: increased resistance to apoptosis enhances the aging process. Cell Mol Life Sci 2011; 68: 1021–1031.

47. Lee JM, Yoshida M, Kim MS, Lee JH, Baek AR, Jang AS, Kim DJ, Minagawa S, Chin SS, Park CS, Kuwano K, Park SW, Araya J. Involvement of Alveolar Epithelial Cell Necroptosis in Idiopathic Pulmonary Fibrosis Pathogenesis. Am J Respir Cell Mol Biol 2018; 59: 215–224.

48. Han J, Zhong CQ, Zhang DW. Programmed necrosis: backup to and competitor with apoptosis in the immune system. Nat Immunol 2011; 12: 1143–1149.

49. Tao P, Sun J, Wu Z, Wang S, Wang J, Li W, Pan H, Bai R, Zhang J, Wang Y, Lee PY, Ying W, Zhou Q, Hou J, Wang W, Sun B, Yang M, Liu D, Fang R, Han H, Yang Z, Huang X, Li H, Deuitch N, Zhang Y, Dissanayake D, Haude K, McWalter K, Roadhouse C, MacKenzie JJ, Laxer RM, Aksentijevich I, Yu X, Wang X, Yuan J, Zhou Q. A dominant autoinflammatory disease caused by non-cleavable variants of RIPK1. Nature 2020; 577: 109–114.

50. Muller T, Dewitz C, Schmitz J, Schroder AS, Brasen JH, Stockwell BR, Murphy JM, Kunzendorf U, Krautwald S. Necroptosis and ferroptosis are alternative cell death pathways that operate in acute kidney failure. Cell Mol Life Sci 2017; 74: 3631–3645.

51. Gautheron J, Vucur M, Schneider AT, Severi I, Roderburg C, Roy S, Bartneck M, Schrammen P, Diaz MB, Ehling J, Gremse F, Heymann F, Koppe C, Lammers T, Kiessling F, Van Best N, Pabst O, Courtois G, Linkermann A, Krautwald S, Neumann UP, Tacke F, Trautwein C, Green DR, Longerich T, Frey N, Luedde M, Bluher M, Herzig S, Heikenwalder M, Luedde T. The necroptosis-inducing kinase RIPK3 dampens adipose tissue inflammation and glucose intolerance. Nat Commun 2016; 7: 11869.

52. Choi ME, Price DR, Ryter SW, Choi AMK. Necroptosis: a crucial pathogenic mediator of human disease. JCI Insight 2019; 4.

53. Rodriguez DA, Weinlich R, Brown S, Guy C, Fitzgerald P, Dillon CP, Oberst A, Quarato G, Low J, Cripps JG, Chen T, Green DR. Characterization of RIPK3-mediated phosphorylation of the activation loop of MLKL during necroptosis. Cell Death Differ 2016; 23: 76–88.

54. McComb S, Shutinoski B, Thurston S, Cessford E, Kumar K, Sad S. Cathepsins limit macrophage necroptosis through cleavage of Rip1 kinase. J Immunol 2014; 192: 5671–5678.

55. Droga-Mazovec G, Bojic L, Petelin A, Ivanova S, Romih R, Repnik U, Salvesen GS, Stoka V, Turk V, Turk B. Cysteine cathepsins trigger caspase-dependent cell death through cleavage of bid and antiapoptotic Bcl-2 homologues. J Biol Chem 2008; 283: 19140–19150.

